# Modulation of the primary auditory thalamus when recognising speech with background noise

**DOI:** 10.1101/510164

**Authors:** Paul Glad Mihai, Nadja Tschentscher, Katharina von Kriegstein

**Affiliations:** Chair of Cognitive and Clinical Neuroscience, Faculty of Psychology, Technische Universität Dresden, 01187 Dresden, Germany; Max Planck Institute for Cognitive and Brain Sciences, 4103 Leipzig, Germany; Research Unit Biological Psychology, Department of Psychology, Ludwig-Maximilians-University Munich, 80802 Munich, Germany

## Abstract

Recognising speech in background noise is a strenuous daily activity, yet most humans can master it. An explanation of how the human brain deals with such sensory uncertainty during speech recognition is to-date missing. Previous work has shown that recognition of speech without background noise involves modulation of the auditory thalamus (medial geniculate body, MGB): There are higher responses in left MGB for speech recognition tasks that require tracking of fast-varying stimulus properties in contrast to relatively constant stimulus properties (e.g., speaker identity tasks) despite the same stimulus input. Here we tested the hypotheses that (i) this task-dependent modulation for speech recognition increases in parallel with the sensory uncertainty in the speech signal, i.e., the amount of background noise and that (ii) this increase is present in the ventral MGB, which corresponds to the primary sensory part of the auditory thalamus. In accordance with our hypothesis, we show—by using ultra-high-resolution functional magnetic resonance imaging in human participants—that the task-dependent modulation of the left vMGB for speech is particularly strong when recognizing speech in noisy listening conditions in contrast to situations where the speech signal is clear. Exploratory analyses showed that this finding was specific to the left vMGB; it was not present in the right vMGB or the midbrain structure of the auditory pathway (left inferior colliculus, IC). The results imply that speech in noise recognition is supported by modifications at the level of the subcortical sensory pathway providing driving input to the auditory cortex.

**Significance Statement:** Speech recognition in noisy environments is a challenging everyday task. One reason why humans can master this task is the recruitment of additional cognitive resources as reflected in recruitment of non-language cerebral cortex areas. Here, we show that also modulation in the primary sensory pathway is specifically involved in speech in noise recognition. We found that the left primary sensory thalamus (ventral medial geniculate body, vMGB) is more involved when recognizing speech signals as opposed to a control task (speaker identity recognition) when heard in background noise vs. when the noise was absent. This finding implies that the brain optimises sensory processing in subcortical sensory pathway structures in a task-specific manner to deal with speech recognition in noisy environments.

## 1. Introduction

Roaring engines, the hammering from a construction site, the chit-chat of many children in a classroom are just some examples of background noises which continuously accompany us. Nevertheless, humans have a remarkable ability to hear and understand the conversation partner, even under these severe listening conditions (Cherry, 1953).

Understanding speech in noise is a complex task that involves both sensory and cognitive processes (Moore et al., 1985; Bregman, 1994; Best et al., 2007; Sayles and Winter, 2008; Shinn-Cunningham and Best, 2008; Song et al., 2010; Adank, 2012; Bronkhorst, 2015; Peelle, 2018; Alavash et al., 2019). However, a more mechanistic explanation of why the human brain masters speech recognition in noise relatively well is missing. Such explanation could advance the understanding of difficulties with speech-in-noise perception in several clinical populations such as age-related hearing impairment (Schoof and Rosen, 2016), autism spectrum disorder (Alcántara et al., 2004), auditory processing disorder (Iliadou et al., 2017), or developmental dyslexia (Chandrasekaran et al., 2009; Ziegler et al., 2009). Furthermore, a more mechanistic understanding of speech-in-noise recognition might also trigger new insight on why artificial speech recognition systems still have difficulties with noisy situations (Scharenborg, 2007; Gupta et al., 2016).

One mechanistic account of brain function that attempts to explain how the human brain deals with uncertainty in the stimulus input is the Bayesian brain hypothesis. It assumes that the brain represents information probabilistically and uses an internal generative model and predictive coding for the most effective processing of sensory input (Knill and Pouget, 2004; Friston, 2005; Kiebel et al., 2008; Friston and Kiebel, 2009). Such type of processing has the potential to explain why the human brain is robust to sensory uncertainty, e.g., when recognising speech despite noise in the speech signal (Srinivasan et al., 1982; Knill and Pouget, 2004). Although predictive coding is often discussed in the context of cerebral cortex organization (Hesselmann et al., 2010; Shipp et al., 2013), it may also be a governing principle of the interactions between cerebral cortex and subcortical sensory pathway structures (Mumford, 1992; von Kriegstein et al., 2008; Huang and Rao, 2011; Bastos et al., 2012; Adams et al., 2013; Seth and Friston, 2016).

In humans, responses in the auditory sensory thalamus (medial geniculate body, MGB) are higher for speech tasks (that emphasise recognition of fast-varying speech properties) in contrast to control tasks (that require recognition of relatively constant properties of the speech signal, such as the speaker identity or the sound intensity level). This response difference holds even if the stimulus input is the same (von Kriegstein et al., 2008; Díaz et al., 2012), indicating that the effect is dependent on the specific tasks. We will therefore call it task-dependent modulation in the following. The task-dependent modulation seems to be behaviourally relevant for speech recognition: performance level in auditory speech recognition positively correlates with the amount of task-dependent modulation in the MGB of the left hemisphere (von Kriegstein et al., 2008; Mihai et al., 2019). This behaviourally relevant task-dependent modulation was located in the ventral part of the MGB (vMGB), which is the primary subsection of the MGB (Mihai et al., 2019). These findings have been interpreted by extending the Bayesian brain hypothesis to cortico-subcortical interactions: cerebral cortex areas provide dynamic predictions about the incoming sensory input to the sensory thalamus to optimally encode the trajectory of the fast-varying and predictable speech input (von Kriegstein et al., 2008; Díaz et al., 2012). If this is the case, then the task-dependent modulation of the vMGB should be especially strong when the fast dynamics of speech have to be recognised in conditions with high sensory uncertainty (Yu and Dayan, 2005; Feldman and Friston, 2010; Díaz et al., 2012; Van de Cruys et al., 2014), for example when the incoming signal is disturbed (Yu and Dayan, 2005; Friston and Kiebel, 2009; Feldman and Friston, 2010; Gordon et al., 2017). In the present study we tested this hypothesis.

## 2. Materials and Methods

### 2.1 Study overview

Presentation of speech in background noise is an ecologically valid way to increase uncertainty about the speech input (Chandrasekaran and Kraus, 2010a). We, therefore, tested, whether the task-dependent modulation of the left vMGB for speech is higher when the speech stimuli are embedded in a noisy as opposed to a clear background. We used ultra-high field functional magnetic resonance imaging (fMRI) at 7 T and a design that has been shown to elicit task-dependent modulation of the MGB in previous studies (von Kriegstein et al., 2008; Díaz et al., 2012). We complemented the design by a noise factor: the speech stimuli (i.e., vowel-consonant-vowel syllables) were presented with and without background noise (Figure 1). The experiment was a 2 × 2 factorial design with the factors task (speech task, speaker task) and noise (noise, clear). To test our hypothesis, we performed a task × noise interaction analysis with the prediction that the task-dependent modulation of the left vMGB increases with decreasing signal-to-noise ratios (i.e., increasing uncertainty about the speech sounds). We focused on the left vMGB for two reasons. First, its response showed behavioural relevance for speech recognition in previous studies (von Kriegstein et al., 2008; Mihai et al., 2019). Second, developmental dyslexia – a condition that is often associated with speech-in-noise recognition difficulties (Chandrasekaran et al., 2009; Ziegler et al., 2009) – has been associated with reduced task-dependent modulation of the left MGB in comparison to controls (Díaz et al., 2012) as well as decreased connections between left MGB and left auditory association cortex (Tschentscher et al., 2019).

**Figure 1.**
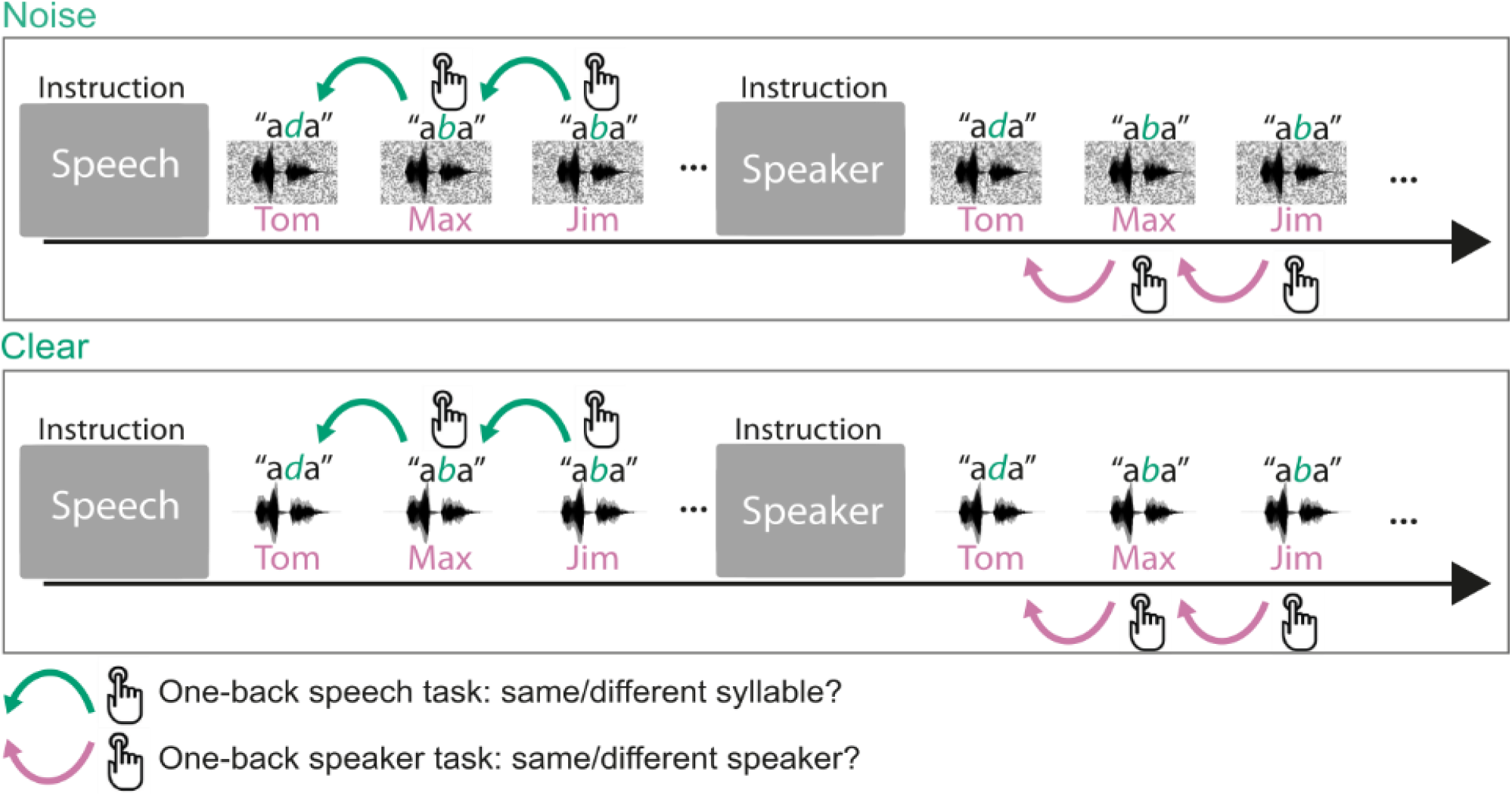
Design and trial structure of the experiment. In the speech task, listeners performed a one-back syllable task. They pressed a button whenever there was a change in syllable in contrast to the immediately preceding one, independent of speaker change. The speaker task used precisely the same stimulus material and trial structure. The task was to press a button when there was a change in speaker identity in contrast to the immediately preceding one, independent of syllable change. The speakers’ voices were resynthesized from the recordings of one speaker’s voice to only differ in constant speaker individuating features (i.e., the vocal tract length and the fundamental frequency of the voice). This ensured that the speaker task could not be done on dynamic speaker individuating features (e.g., idiosyncrasies in pronunciations of phonemes). An initial task instruction screen informed participants about which task to perform. Participants heard stimuli either with concomitant speech-shaped noise (noise condition) or without background noise (clear condition). Thus the experiment had four conditions: speech task/noise, speaker task/noise, speech task/clear, speaker task/clear. Stimuli in the speech and speaker tasks were precisely identical.

In addition to testing our main hypothesis, the design also allowed the exploration of the role of the inferior colliculus (IC) – the midbrain station of the auditory sensory pathway – in speech-in-noise recognition.

### 2.2 Participants

The Ethics committee of the Medical Faculty, University of Leipzig, Germany, approved the study. We recruited 17 participants (mean age 27.7, SD 2.5 years, 10 female; 15 of these participated in a previous study: Mihai et al., 2019) from the database of the Max Planck Institute for Human Cognitive and Brain Sciences (MPI-CBS), Leipzig, Germany. The sample size was based on the amount of data acquisition time allocated by the MPI-CBS directorial board to the study. The participants were right-handed (as assessed by the Edinburgh Handedness Inventory (Oldfield 1971)), and native German speakers. Participants provided written informed consent. None of the participants reported a history of psychiatric or neurological disorders, hearing difficulties, or current use of psychoactive medications. Normal hearing abilities were confirmed with pure tone audiometry (250 Hz to 8000 Hz; Madsen Micromate 304, GN Otometrics, Denmark) with a threshold equal to and below 25 dB). To exclude possible undiagnosed developmental dyslexics, we tested the participant’s reading speed and reading comprehension using the German LGVT: 6-12 test (Schneider et al., 2007). The cut-off for both reading scores was set to those levels mentioned in the test instructions as the “lower average and above” performance range (i.e., 26% - 100% of the calculated population distribution). None of the participants performed below the cut off performance (mean 68.7%, SD 20.6%, lowest mean score: 36%). In addition, participants were tested on rapid automatized naming (RAN) of letters, numbers, and objects (Denckla and Rudel, 1976). The time required to name letters and numbers predicts reading ability and is longer in developmental dyslexics compared with typical readers, whereas the time to name objects is not a reliable predictor of reading ability in adults (Semrud-Clikeman et al., 2000). Participants scored well within the range of control participants for letters (mean 17.25, SD 2.52 s), numbers (mean 16.79, SD 2.63 s), and objects (mean 29.65, SD 4.47 s), based on results from a previous study (Díaz et al., 2012, letters: 16.09, SD 2.60; numbers: 16.49, SD 2.35; objects: 30.84, SD 5.85; age of participants was also comparable 23.5, SD 2.8 years). Furthermore, none of the participants exhibited a clinically relevant number of traits associated with autism spectrum disorder as assessed by the Autism Spectrum Quotient [AQ; mean: 15.9, SD 4.1; cut-off: 32-50; (Baron-Cohen et al., 2001)]. We tested AQ as autism can be associated with difficulties in speech-in-noise perception (Alcántara et al., 2004; Groen et al., 2009). Participants received monetary compensation for participating in the study.

### 2.2 Stimuli

We recorded 79 different vowel-consonant-vowel (VCV) syllables with an average duration of 784 ms, SD 67 ms. These recordings constitute a subsample from those used in (Mihai et al., 2019). These were spoken by one male voice (age 29 years), recorded with a video camera (Canon Legria HFS10, Canon, Japan) and a Røde NTG-1 microphone (Røde Microphones, Silverwater, NSW, Australia) connected to a pre-amplifier (TubeMP Project Series, Applied Research and Technology, Rochester, NY, USA) in a sound-attenuated room. The sampling rate was 48 kHz at 16 bit. Auditory stimuli were cut and flanked by Hamming windows of 15 ms at the beginning and end, converted to mono, and root-mean-square equalised using Python 3.6 (Python Software Foundation, www.python.org). The 79 auditory files were resynthesized with TANDEM-STRAIGHT (Banno et al., 2007) to create three different speakers: 79 auditory files with a vocal tract length (VTL) of 17 cm and glottal pulse rate (GPR) of 100 Hz, 79 with VTL of 16 cm and GPR of 150 Hz, and 79 with VTL of 14 cm and GPR of 300 Hz. This procedure resulted in 237 different auditory stimuli. The parameter choice (VTL and GPR) was motivated by the fact that a VTL difference of 25% and a GPR difference of 45% suffices for listeners to hear different speaker identities (Gaudrain et al., 2009a; Kreitewolf et al., 2014). Additionally, we conducted pilot experiments (12 pilot participants which did not participate in the main experiment) in order to fine-tune the combination of VTL and GPR that resulted in a balanced behavioural accuracy score between the speech and speaker tasks. The pilot experiments were conducted outside the MRI-machine, but included continuous recordings of MRI-gradient noise to simulate a real MRI-environment.

We embedded the 237 stimuli in background noise to create the stimuli for the condition with background noise. The background noise consisted of normally distributed random (white) noise filtered with a speech-shaped envelope. We calculated the envelope from the sum of all VCV stimuli presented in the experiment. We used speech-shaped noise as it has a stronger masking effect than stationary random non-speech noise (Carhart et al., 1975). Before each experimental run, the noise was computed and added to the stimuli included in the run with a signal-to-noise ratio (SNR) of 2 dB. The SNR choice was based on a pilot study that showed a performance decrease of at least 5% but no greater than 15% between the clear and noise condition. In the pilot study, we started at an SNR of −10 dB and increased this value until we converged on an SNR of 2 dB. Calculations were performed in Matlab 8.6 (The Mathworks Inc., Natick, MA, USA) on Ubuntu Linux 16.04 (Canonical Ltd., London, UK).

### 2.3 Procedure

We conceived the experiment as a 2 × 2 factorial design. The first factor was task (speech, speaker) similar to previous experiments that reported task-dependent modulation of the MGB (von Kriegstein et al., 2008; Díaz et al., 2012; Mihai et al., 2019). The second factor was background noise (clear, noise, Figure 1). Participants listened to blocks of auditory VCV syllables and were asked to perform the two types of tasks: the speech task and the speaker task. In the speech task, participants reported via button press whether the current syllable was different from the previous one (1-back task). In the speaker task, participants reported via button press whether the current speaker was different from the previous one. The blocks had either syllables with background noise (noise condition) or without background noise (clear condition).

Task instructions were presented for two seconds before each block and consisted of white written words on a black background (German words “Silbe” for syllable indicating the speech task, and “Person” for person indicating the speaker task). After the instruction, the block of syllables started (Figure 1). Each block contained twelve stimuli. Each stimulus had a duration of approximately 784 ms, and the stimulus presentation was followed by 400 ms of silence. Within one block both syllables and speakers changed at least twice, with a theoretical maximum of nine changes. The theoretical maximum was derived from random sampling of seven instances from three possible change types: no change, speech change, speaker change, and change of speech and speaker. The average length of a block was 15.80 seconds, SD 0.52 seconds. The presentation of the stimuli was randomized and balanced with regard to the amount of speaker identity and syllable changes within a block. The same block containing speaker identity changes also contained syllable changes. These blocks were repeated, once with the instruction to perform the speaker identity task and the other time to perform the speech task. This procedure ensured that subjects heard exactly the same stimuli while performing the two different tasks.

The experiment was divided into four runs. The first three runs had a duration of 12:56 min and included 40 blocks: 10 for each of the four conditions (speech task/noise, speaker task/noise, speech task/clear, speaker task/clear). A fourth run had a duration of 6:32 min and included 20 blocks (5 for each of the four conditions). For two participants, only the first three runs were recorded due to time constraints. Participants could rest for one minute between runs.

Participants were familiarised with the three speakers’ voices to ensure that they could perform the speaker-identity task of the main experiment. The speaker familiarisation took place 30 minutes before the fMRI experiment. It consisted of a presentation of the speakers and a test phase. In the presentation phase, the speakers were presented in six blocks, each containing nine pseudo-randomly chosen VCV stimuli from the 237 total. Each block contained one speaker-identity only. Participants were alerted to the onset of a new speaker identity block by the presentation of white words on a black screen indicating speaker 1, speaker 2, or speaker 3. Participants listened to the voices with the instruction to memorise the speaker’s voice. In the following test phase participants were presented with four blocks of nine trials that each contained randomly chosen syllable pairs spoken by the three speakers. The syllable pairs could be from the same or a different speaker. We asked participants to indicate whether the speakers of the two syllables were the same by pressing keypad buttons “1” for yes and “2” for no. Participants received visual feedback for correct (the green flashing German word for correct: “Richtig”) and incorrect (the red flashing German word for incorrect: “Falsch”) answers. The speaker familiarisation consisted of three 2:50 min runs (each run contained one presentation and one test phase). If participants scored below 80% on the last run, they performed an additional run until they scored above 80%. All participants exceeded the 80% cut-off value.

The experiments were programmed in the Matlab Psychophysics Toolbox [Psychtoolbox-3, www.psychtoolbox.com (Brainard, 1997)] running on Matlab 8.6 (The Mathworks Inc., Natick, MA, USA) on Ubuntu Linux 16.04 (Canonical Ltd., London, UK). The sound was delivered through a MrConfon amplifier and headphones (manufactured 2008; MrConfon GmbH, Magdeburg, Germany).

### 2.4 Data Acquisition and Processing

MRI data were acquired using a Siemens Magnetom 7 T scanner (Siemens AG, Erlangen, Germany) with an 8-channel head coil. We convened on the 8-channel coil, due to its spaciousness which allowed the use of higher quality headphones (manufactured 2008; MrConfon GmbH, Magdeburg, Germany). Functional MRI data were acquired using echo-planar imaging (EPI) sequences. We used partial brain coverage with 30 slices. The volume was oriented in parallel to the superior temporal gyrus such that the slices encompassed the MGB, the inferior colliculi (IC), and the Heschl’s gyrus.

The EPI sequences had the following acquisition parameters: TR = 1600 ms, TE = 19 ms, flip angle 65°, GRAPPA (Griswold et al., 2002) with acceleration factor 2, 33% phase oversampling, matrix size 88, field of view (FoV) of 132 mm x 132 mm, phase partial Fourier 6/8, voxel size 1.5 mm isotropic resolution, interleaved acquisition, anterior to posterior phase-encode direction. The first three runs consisted of 485 volumes (12:56 min), and the fourth run consisted of 245 volumes (6:32 min). During functional MRI data acquisition, we also acquired physiological values (heart rate, and respiration rate) using a BIOPAC MP150 system (BIOPAC Systems Inc., Goleta, CA, USA).

To address geometric distortions in EPI images we recorded gradient echo based field maps which had the following acquisition parameters: TR = 1500 ms, TE1 = 6.00 ms, TE2 = 7.02 ms, flip angle 60°, 0% phase oversampling, matrix size 100, FoV 220 mm x 220 mm, phase partial Fourier off, voxel size 2.2 mm isotropic resolution, interleaved acquisition, anterior to posterior phase-encode direction. Resulting images from field map recordings were two magnitude images and one phase difference image.

Structural images were recorded using an MP2RAGE (Marques et al., 2010) T1 protocol: 700 μm isotropic resolution, TE = 2.45ms, TR = 5000 ms, TI1 = 900 ms, TI2 = 2750 ms, flip angle 1 = 5°, flip angle 2 = 3°, FoV 224 mm × 224 mm, GRAPPA acceleration factor 2, duration 10:57 min.

### 2.5 Behavioural Data Analysis

Button presses (hits, misses) were binomially distributed, and were thus modeled using a binomial logistic regression which predicts the probability of correct button presses based on four independent variables (speech task/noise, speaker task/noise, speech task/clear, speaker task/clear) in a Bayesian framework (McElreath, 2018).

To pool over participants and runs we modelled the correlation between intercepts and slopes. For the model implementation and data analysis, we used PyMC3 3.5 (Salvatier et al., 2016), a probabilistic programming package for Python 3.6. We sampled with a No-U-Turn Sampler (Hoffman and Gelman, 2014) with four parallel chains. Per chain, we had 5,000 samples with 5,000 as warm-up. The data entering the model was mean centered by subtracting the mean and dividing by two standard deviations (Gelman and Hill, 2006). This transformation does not change the fit of the linear model and the coefficients are interpretable in comparison to the mean of the data. The reason behind this transformation is the faster and more accurate convergence of the Markov Chain sampling (McElreath, 2018).

There were the following effects of interest: main effects (clear - noise, speech task - speaker task), the interaction (speech task/ noise - speaker task/ noise) - (speech task/ clear - speaker task/ clear), simple main effects (speech task/ noise - speaker task/ noise, speech task/ clear - speaker task/ clear). For the effects of interest, we calculated means from the posterior distributions and 95% highest posterior density intervals (HDP). The HPD is the probability that the mean lies within the interval (Gelman et al., 2013; McElreath, 2018), this means that we are 95% sure the mean lies within the specified interval bounds. If the posterior probability distribution of odds ratios does not strongly overlap one (i.e., the HPD excludes one), then it is assumed that there is a detectable difference between conditions (Bunce and McElreath, 2017; McElreath, 2018).

The predictors included in the behavioural data model were: task (*XS*:1 = speech task, 0 = speaker task), and background noise (*XN*: 1 = noise, 0 = clear). We also included the two-way interaction of task and noise condition. Because data were collected across participants and runs, we included random effects for both of these in the logistic model. Furthermore, since ~11% of the data exhibited ceiling effects (i.e., some participants scored at the highest possible level) which would result in underestimated means and standard deviations (Uttl, 2005), we treated these data as right-censored and modeled them using a Potential class (Lauritzen et al., 1990; Jordan, 1998) as implemented in PyMC3. This method integrates the censored values using the log of the complementary normal cumulative distribution function (Gelman et al., 2013; McElreath, 2018). In essence, we sampled twice, once for the observed values without the censored data points, and once for the censored values only. The model is described below.

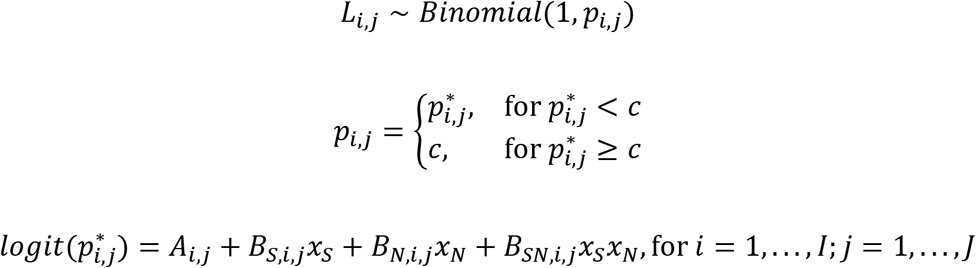

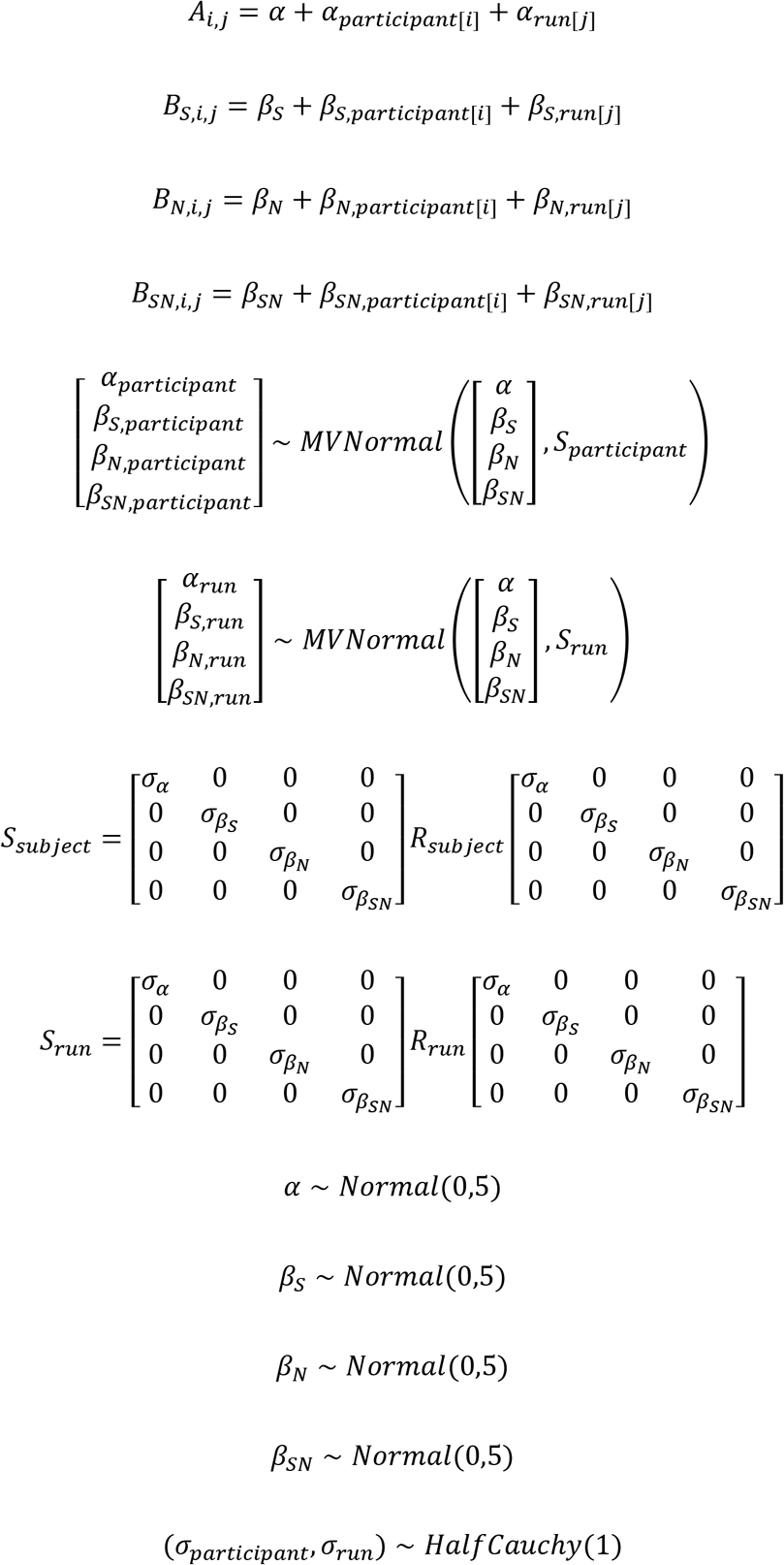

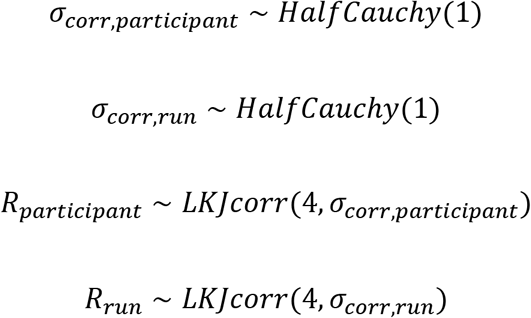

*I* represents the participants and *J* the runs. The model is compartmentalized into sub-models for the intercepts and slopes. *A_i,j_* is the sub-model for the intercept for observations *i,j.* Similarly, *B_S,i,j_, B_N,i,j_*, and *B_SN,i,j_* are the sub-models for the speech task - speaker task slope, clear-noise slope and the interaction slope, respectively; *S_subject_/S_run_* are the covariance matrices for participant/run. *R_subject_/B_run_* are the priors for the correlation matrices modelled as LKJ probability densities (Lewandowski et al., 2009). Weakly informative priors for the intercept (*α*) and additional coefficients (e.g., *β_s_*), random effects for participant and run (*β_S,subject_, β_S,run_*), and multivariate priors for participants and runs identify the model by constraining the position of *p_ij_* to reasonable values. Here we used normal distributions as priors. Furthermore, *p_t,j_* is defined as the ramp function equal to the proportion of hits when these are known and below the ceiling (*c*), and set to the ceiling if they are equal to or greater than the ceiling *c*.

We additionally analyzed the reaction times, similarly to the model described above but without consideration of ceiling effects as they are non-existent. Posterior distributions were computed for each condition, and we computed main effects and the interaction between task and noise. If the posterior probability distribution of the difference scores and the interaction does not strongly overlap zero (i.e., the HPD excludes zero), then it is assumed that there is a detectable difference (Bunce and McElreath, 2017; McElreath, 2018).

### 2.6 Functional MRI Data Analysis

#### 2.6.1 Preprocessing of fMRI data

The MP2RAGE images were first segmented using SPM’s segment function (SPM 12, version 12.6906, Wellcome Trust Centre for Human Neuroimaging, UCL, UK, http://www.fil.ion.ucl.ac.uk/spm) running on Matlab 8.6 (The Mathworks Inc., Natick, MA, USA) in Ubuntu Linux 16.04 (Canonical Ltd., London, UK). The resulting grey and white matter segmentations were summed and binarised to remove voxels that contain air, scalp, skull and cerebrospinal fluid from structural images using the ImCalc function of SPM.

We used the template image created for a previous study (Mihai et al., 2019) using structural MP2RAGE images from the 28 participants of that study. We chose this template since 15 participants in the current study are included in this image, and the vMGB mask (described below) is in the same space as the template image. The choice of this common template reduces warping artefacts, which would be introduced with a different template, as both the vMGB mask and the functional data of the present study would need to be warped to a common space. The template was created and registered to MNI space with ANTs (Avants et al., 2008) and the MNI152 template provided by FSL 5.0.8 (Smith et al., 2004). All MP2RAGE images were preprocessed with Freesurfer (Fischl et al., 2004; Han and Fischl, 2007) using the recon-all command to obtain boundaries between grey and white matter, which were later used in the functional to structural registration step.

Preprocessing and statistical analyses pipelines were coded in nipype 1.1.2 (Gorgolewski et al., 2011). Head motion and susceptibility distortion by movement interaction of functional runs were corrected using the Realign and Unwarp method (Andersson et al., 2001) in SPM 12. This step also makes use of a voxel displacement map (VDM), which addresses the problem of geometric distortions in EPI caused by magnetic field inhomogeneity. The VDM was calculated using field map recordings, which provided the absolute value and the phase difference image files, using the FieldMap Toolbox (Jezzard and Balaban, 1995) of SPM 12. Outlier runs were detected using ArtifactDetect (composite threshold of translation and rotation: 1; intensity Z-threshold: 3; global threshold: 8; https://www.nitrc.org/projects/artifact_detect/). Coregistration matrices for realigned functional runs per participant were computed based on each participant’s structural image using Freesurfer’s BBregister function (register mean EPI image to T1). We used a whole-brain EPI volume as an intermediate file in the coregistration step to avoid registration problems due to the limited FoV of the functional runs. Warping using coregistration matrices (after conversion to the ITK coordinate system) and resampling to 1 mm isovoxel was performed using ANTs. Before model creation, we smoothed the data in SPM12 using a 1 mm kernel at full-width half-maximum.

#### 2.6.2 Physiological data

Physiological data (heart rate and respiration rate) were processed by the PhysIO Toolbox (Kasper et al., 2017) to obtain Fourier expansions of each, in order to enter these into the design matrix (see section 2.6.3 Testing our hypothesis in the left vMGB). Since heartbeats and respiration result in undesired cortical and subcortical artefacts, regressing these out increases the specificity of fMRI responses to the task of interest (Kasper et al., 2017). These artefacts occur in abundance around the thalamus (Kasper et al., 2017).

#### 2.6.3 Testing our hypothesis in the left vMGB

Models were set up in SPM 12 using the native space data for each participant. We modelled five conditions of interest: speech task/noise, speaker task/noise, speech task/clear, speaker task/clear, and task instruction. Onset times and durations were used to create boxcar functions, which were convolved with the hemodynamic response function (HRF) provided by SPM 12. The design matrix also included the following nuisance regressors: three cardiac, four respiratory, and a cardiac × respiratory interaction regressor. We additionally entered the outlier regressors from the ArtifactDetect step.

Parameter estimates were computed for each condition at the first level using restricted maximum likelihood (REML) as implemented in SPM 12. Parameter estimates for each of the four conditions of interest (speech task/noise, speaker task/noise, speech task/clear, speaker task/clear) were registered to the MNI structural template using a two-step registration in ANTs. First, a quick registration was performed on the whole head using rigid, affine and diffeomorphic transformations (using Symmetric Normalization, SyN), and the mutual information similarity metric. Second, the high-quality registration was confined to the volume that was covered by the 30 slices of the EPI images. These volumes include the IC, MGB, and primary and secondary auditory cortices. This step used affine and SyN transformations and mean squares and neighbourhood cross-correlation similarity measures. We performed the registration to MNI space by linearly interpolating the contrast images using the composite transforms from the high-quality registration.

We extracted parameter estimates for each of the four conditions of interest per participant, averaged over all voxels from the region of interest, i.e., the left vMGB. To locate the left vMGB, we used the mask from (Mihai et al., 2019), which included 15 of the 17 participants of the present study (Figure 2).

**Figure 2.**
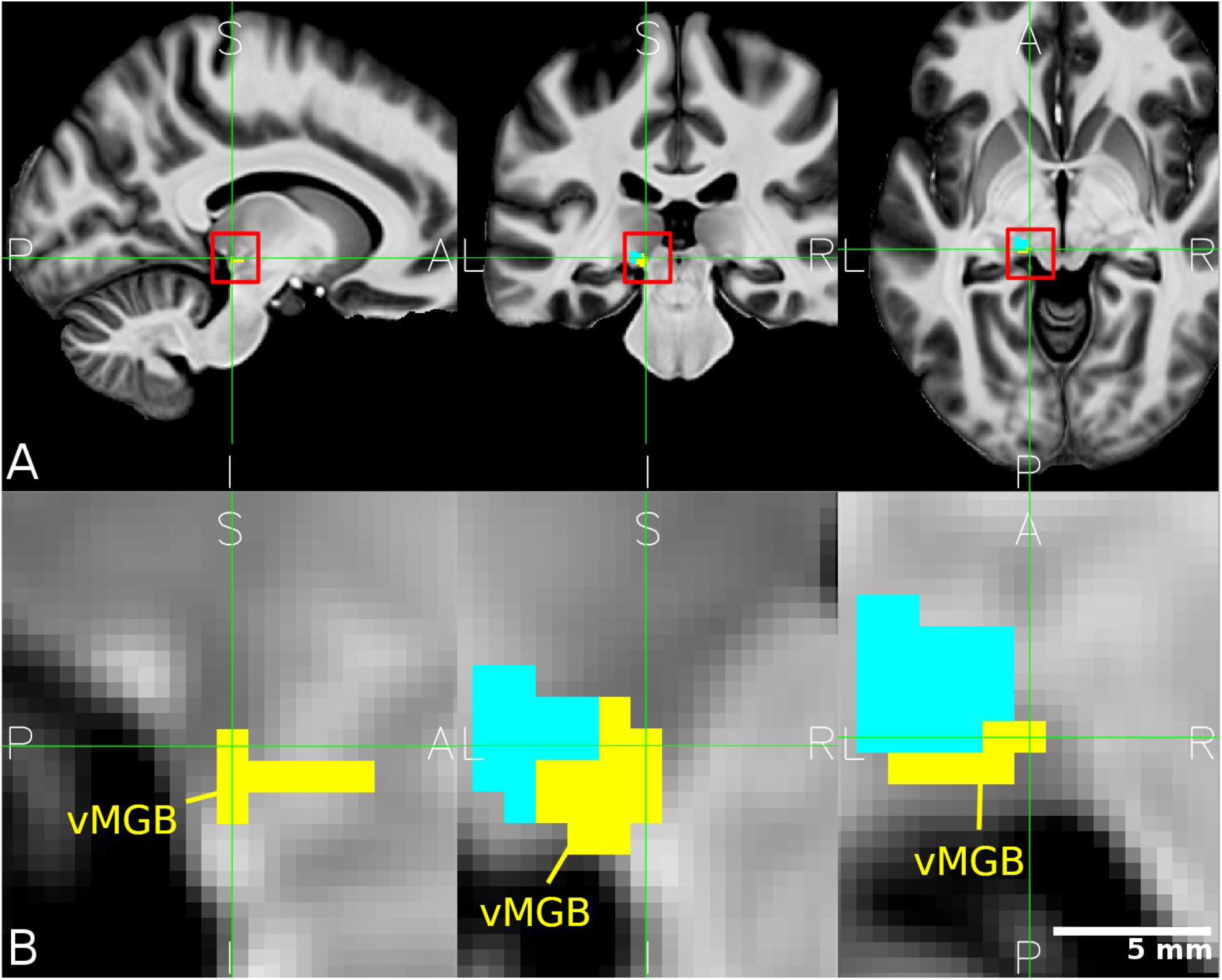
Location of the left MGB masks. (A) The mean structural image across participants (n = 33) in MNI space. The red squares denote the approximate location of the left MGB and encompass the zoomed in view in B. (B) Closeup of the left vMGB (yellow). The tonotopic gradient two is shown in cyan. Panels correspond to sagittal, coronal, and axial slices (P: posterior, A: anterior, S: superior, I: inferior, L: left, R: right).

We analysed the extracted parameter estimates in a Bayesian framework (McElreath, 2018). The data entering the model was mean centered by subtracting the mean and dividing by two standard deviations (Gelman and Hill, 2006). This transformation does not change the fit of the linear model and the coefficients are interpretable in comparison to the mean of the data. The reason behind this transformation is the faster and more accurate convergence of the Markov Chain sampling (McElreath, 2018). The model was implemented in PyMC3 with a No-U-Turn Sampler with four parallel chains. Per chain, we sampled posterior distributions which had 5000 samples with 5000 as warm-up. The predictors included in the model were: task (*XS*: 1 = speech task, 0 = speaker task), and background noise (*XN*: 1 = noise, 0 = clear). We also included the two-way interaction of task and noise condition. Because data were collected across participants, it was reasonable to include random effects. To pool over participants, we modelled the correlation between intercepts and slopes over participants. The interaction model is described below.

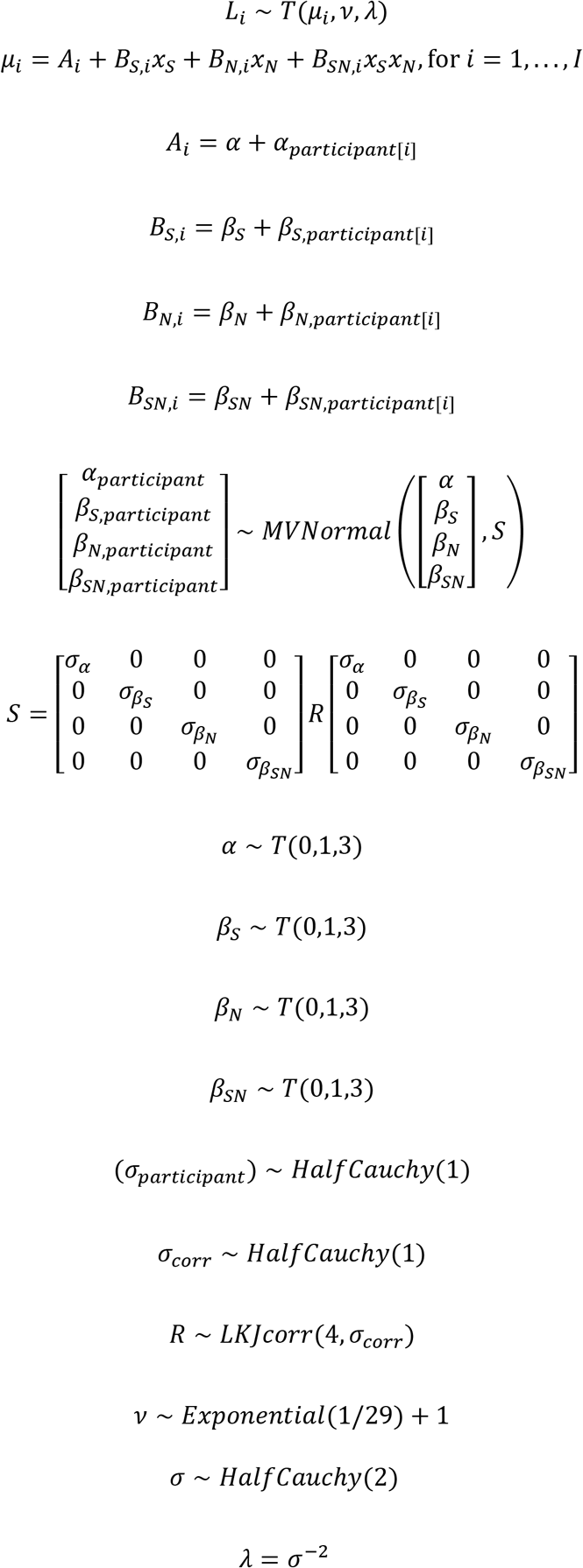

*I* represents the participants. The model is compartmentalized into sub-models for the intercepts and slopes. *A_i_* is the sub-model for the intercept for observations *i.* Similarly, *B_S,i_, B_N,i_*, and *B_SN,i_* are the sub-models for the speech task - speaker task slope, clear-noise slope and the interaction slope, respectively; S is the covariance matrix and *R* is the prior for the correlation matrix modelled as an LKJ probability density (Lewandowski et al., 2009). Weakly informative priors for the intercept (*α*) and additional coefficients (e.g., *β_S_*), random effects for participant (*β_S,subject_*), and multivariate priors for participants identify the model by constraining the position of *μ,* to reasonable values. Here we used Student’s*-T* distributions as priors.

From the model output, we calculated posterior distributions for each condition of interest (speech task/noise, speaker task/ noise, speech task/clear, speaker task/clear). Posterior distributions, in comparison to point estimates, have the advantage of quantifying uncertainty about each parameter. We summarised each posterior distribution using the mean as a point estimate (posterior mean) together with a 95% highest posterior density interval (HPD). The HPD is the probability that the mean lies within the interval (Gelman et al., 2013; McElreath, 2018), e.g., we are 95% sure the mean lies within the specified interval bounds. We computed the following contrasts of interest: interaction (speech task/noise - speaker task/noise) – (speech task/clear – speaker task/clear); simple main effects (speech task/noise – speaker task/noise), (speech task/clear – speaker task/clear); main effect of task (speech task – speaker task). Differences between conditions were converted to effect sizes [Hedges g* (Hedges and Olkin, 1985)]. Hedges g*, like Cohen’s d (Cohen, 1988), is a population parameter that computes the difference in means between two variables normalised by the pooled standard deviation with the benefit of correcting for small sample sizes. Based on Cohen (1988), we interpreted effect sizes on a spectrum ranging from small (g* ≈ 0.2), to medium (g* ≈ 0.5), to large (g* ≈ 0.8), and beyond. If the HPD did not overlap zero, we considered this to be a robust effect (Bunce and McElreath, 2017; McElreath, 2018). However, we caution readers that if the HPD includes zero, it does not mean that the effect is missing (Amrhein et al., 2019). Instead, we quantify and interpret the magnitude (by the point estimate) and its uncertainty (by the HPD) provided by the data and our assumptions (Anderson, 2019).

#### 2.6.4 Analyses of the left inferior colliculus

The study design and acquisition parameters also allowed us to explore the involvement of the IC in speech-in-noise recognition (for a rationale of these exploratory analyses see results, section 3.2.2). To analyse the task × noise interaction and the main effect of task in the bilateral IC we used the same analysis procedures as described for the left vMGB (see section 2.6.3 Testing our hypothesis in the left vMGB). As region of interest, we used the IC masks described in (Mihai et al., 2019) and limited them to the tonotopic parts of the IC, i.e., the central nucleus (Figure 3), which corresponds to the primary auditory pathway (Davis, 2005). We will call it cIC in the following. Furthermore, we performed a Pearson’s correlation calculation to analyse the correlation (speech - speaker task correlated with speech accuracy score) in the left cIC. The motivation for this test was based on similar correlations (i.e., speech – control task correlated with speech accuracy score) found in two previous experiments in the left cIC (von Kriegstein et al., 2008 experiment 1 and 2) (for further details see results, section 3.2.2).

**Figure 3.**
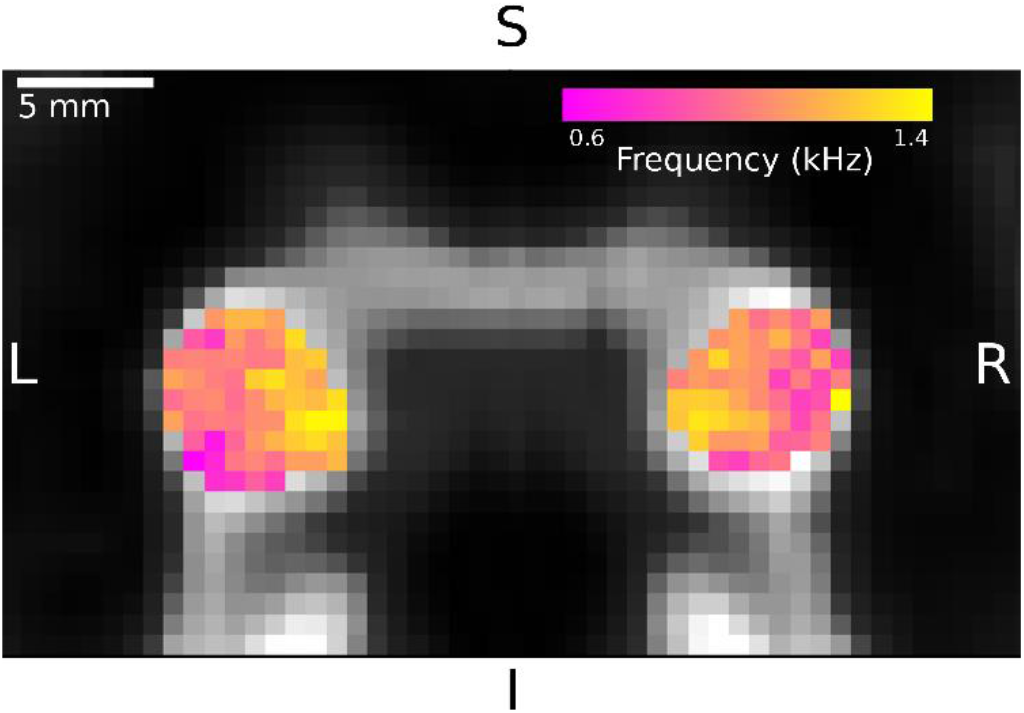
Tonotopy gradients in the inferior colliculi. The colored parts show one slice of the mean tonotopic map across participants in the left and right IC in coronal view (S: superior, I: inferior, L: left, R: right). Individual tonotopies showed high varuability (results not shown). The mean tonotopy revealed a gradient from low frequencies in lateral locations to high frequencies in medial locations (Mihai et al., 2019). The maps were used to construct a region of interest for the central nucleus of the IC (cIC).

### 3. Results

#### 3.1 Behavioural results

##### 3.1.1 Accuracy

Participants performed well above chance level in all four conditions (> 82% correct; Table 1; Figure 4A).

**Figure 4.**
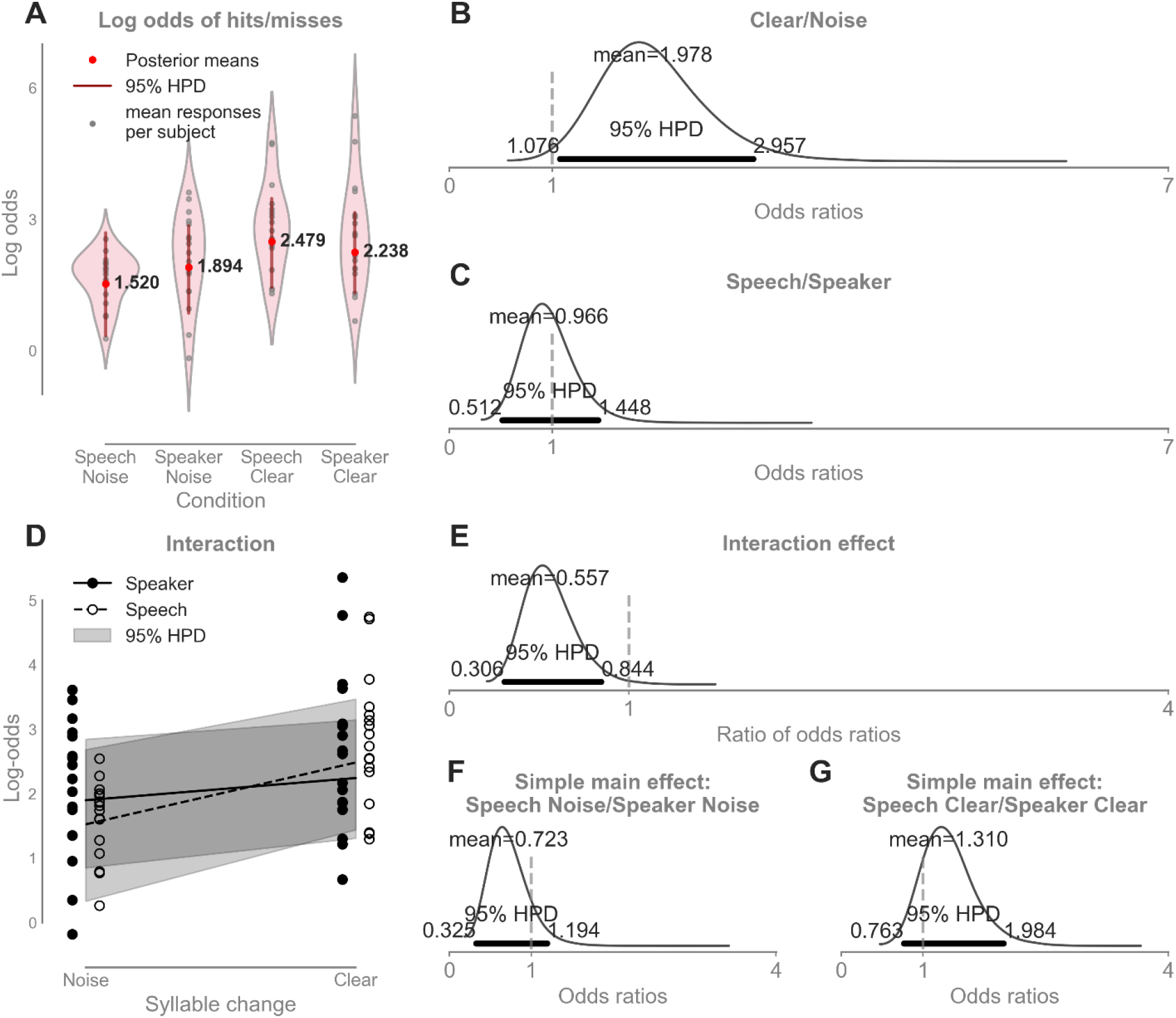
Behavioural results. We performed a binomial logistic regression to compute the rate of hits and misses in each condition because behavioural data were binomially distributed. For this reason, results are reported in log odds and odds ratios. The results showed a detectable main effect of noise and interaction between noise and task. There was no main effect of task, and no detectable simple main effects (speech task/noise - speaker task/noise; speech task/clear - speaker task/clear). **A.** Log odds of hits and misses for each condition. The grey dots indicate mean responses for individual participants, the red dots and accompanying numbers denote the posterior mean per condition, and the dark red lines demarcate the 95% highest posterior density interval (HPD). The rate of hits compared to misses is plotted on a log scale to allow for a linear representation. **B.** Mean odds ratio for the clear and noise conditions. The odds of hits in the clear condition were on average twice as high as in the noise condition (the mean odds ratio was 1.978 [1.076, 2.957]). The HPD excluded 1 and indicated a detectable difference between conditions: No difference would be assumed if the odds ratio was 1 (50/50 chance or 1:1 ratio; Chen, 2003). **C.** Mean odds ratio for the speech task - speaker task conditions. The mean odds ratio was ~1 indicating no difference between the speech and speaker task conditions. **D**. Visualization of the interaction (task × noise) as a comparison of slopes with 95% HPD. **E**. The ratio of odds ratios of the simple main effects speech task/noise - speaker task/noise and speech task/clear - speaker task/clear. The mean and 95% HPD was 0.557 [0.306, 0.844]. The HPD excluded 1 indicating an interaction effect. **F**. Mean odds ratio for the simple main effect speech task/noise - speaker task/noise. The rate of hits in the speech task/noise condition was on average ~1/3 lower than the rate of hits in the speaker task/noise condition; however, the HPD strongly overlapped 1 indicating that there was no difference between conditions. **G**. Mean odds ratio for the simple main effect speech task/clear - speaker task/clear. The rate of hits in the speech task/clear condition was on average ~1/3 higher than the rate of hits in the speaker task/clear condition; however, the HPD strongly overlapped 1 indicating that there was no detectable difference between conditions.

**Table 1.** The proportion of hits for each of the four conditions in the experiment. HDP: highest posterior density interval.

|  | Speech task/<br>Noise | Speaker task/<br>Noise | Speech task/<br>Clear | Speaker task/<br>Clear |
| --- | --- | --- | --- | --- |
| Hit rate<br>[95% HPD] | 0.82 [0.62, 0.95] | 0.87 [0.74, 0.96] | 0.92 [0.83, 0.98] | 0.90 [0.81, 0.97] |

Performing the tasks with background noise was more difficult than the conditions without background noise for both the speech and the speaker task (Figure 4B, for details on statistics, see figure and legend). The rate of hits in the speech task was the same as in the speaker task (Figure 4C). There was a detectable interaction between task and noise (Figure 4D/E), but simple main effects (i.e., speech task/noise - speaker task/noise (Figure 4F) and speech task/clear - speaker task/clear (Figure 4G)) were not present. We also observed ceiling effects in 11% of the cases, which were modeled accordingly (Materials and Methods, section 2.5).

##### 3.1.2 Reaction times

The reaction times analysis showed that for the speech task participants required on average 0.166 [0.114, 0.222] s longer to react than for the speaker task (Figure 5). This effect is explained by the fact that VCV syllables had constant vowels and only the consonants changed within one block. Therefore, listeners had to wait for the consonant to detect a change. Whereas, for the speaker identitiy task the glottal pulse rate is the strongest cue, and is immediately decoded (Gaudrain et al., 2009b). The difference in reaction times between the noise and clear condition was on average 0.059 [0.010, 0.113] s. This difference showed that the noise condition required a minimal amount of extra processing time, yet this difference was on average very small. Lastly, the task x noise interaction was on average 0.022 s with the HPD overlapping zero ([−0.028, 0.076] s), which is not a meaningful effect.

**Figure 5.**
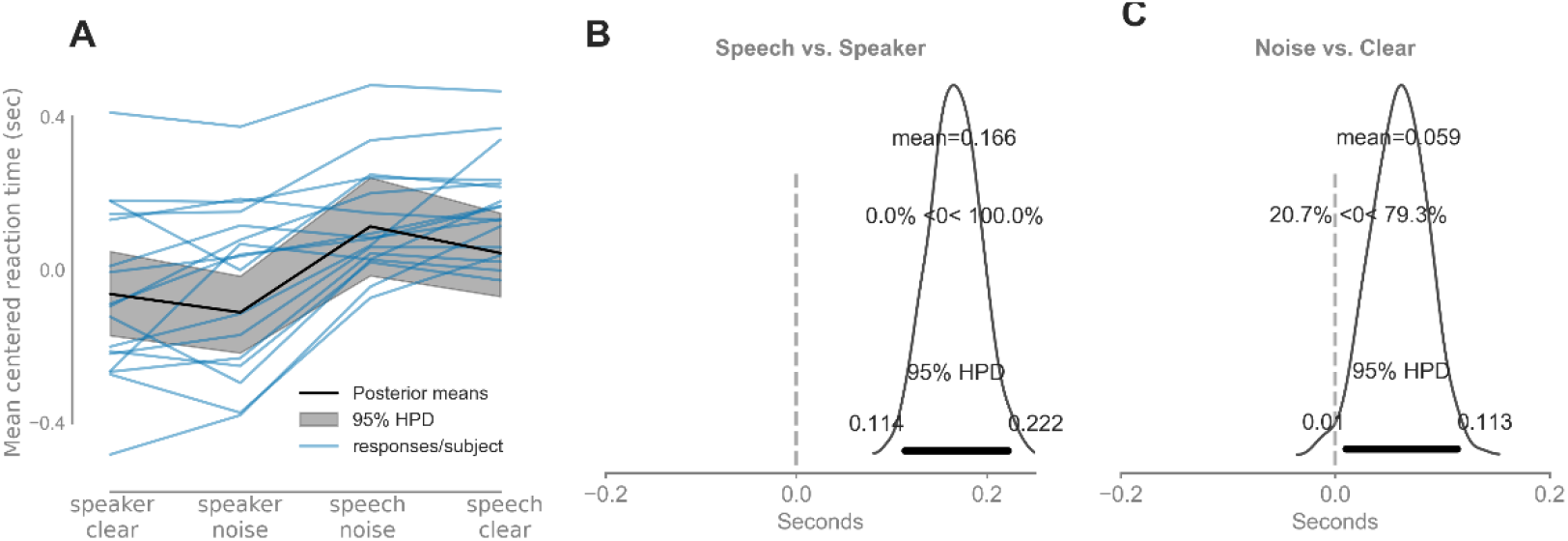
Reaction times results. **A.** Mean centered reaction times for each condition. The blue lines indicate individual average reaction times, the black line denotes the estimated reaction time per condition averaged over participants and runs, the grey shaded area denotes the 95% highest posterior density interval (HPD). **B.** Mean reaction time difference between the Speech and Speaker task. On average, participants took 0.166 [0.114, 0.222] s longer to react in the Speech than to the Speaker task. **C.** Mean reaction time difference between the Noise and the Clear condition. On average, participants took 0.059 [0.010, 0.113] s longer to react during the Noise vs. Clear condition. There was no task x noise interaction.

#### 3.2 fMRI Results

##### 3.2.1 The task-dependent modulation of left vMGB was increased for recognizing speech-in-noise in contrast to the clear speech condition

We localised the left vMGB based on an independent functional localizer (Figure 6B). Following our hypothesis, there was increased BOLD response for the task × noise interaction [(speech task/noise - speaker task/noise) - (speech task/clear - speaker task/clear)] in the left vMGB (Figure 6A/B). The interaction effect had a mean large effect size ranging across participants from a small effect to a very large effect (g*=2.549 [0.211, 5.066]; Figure 6C and D). The 95% HPD of the interaction effect excluded 0, indicating that this was a robust effect (Bunce and McElreath, 2017; McElreath, 2018). Simple main effect analyses showed that the direction of the interaction was as expected. The speech task/noise condition yielded higher left vMGB responses in contrast to the speaker task/noise condition, ranging from a medium to a very large effect across participants (g* = 1.104 [0.407, 1.798]; Figure 6E). Conversely, the left vMGB response difference between the speech task and speaker task in the clear condition had a small effect size (g* = 0.243 [−0.366, 0.854]; Figure 6F), ranging from a negative medium effect to a positive large effect across participants, and the HPD overlapped 0.

**Figure 6.**
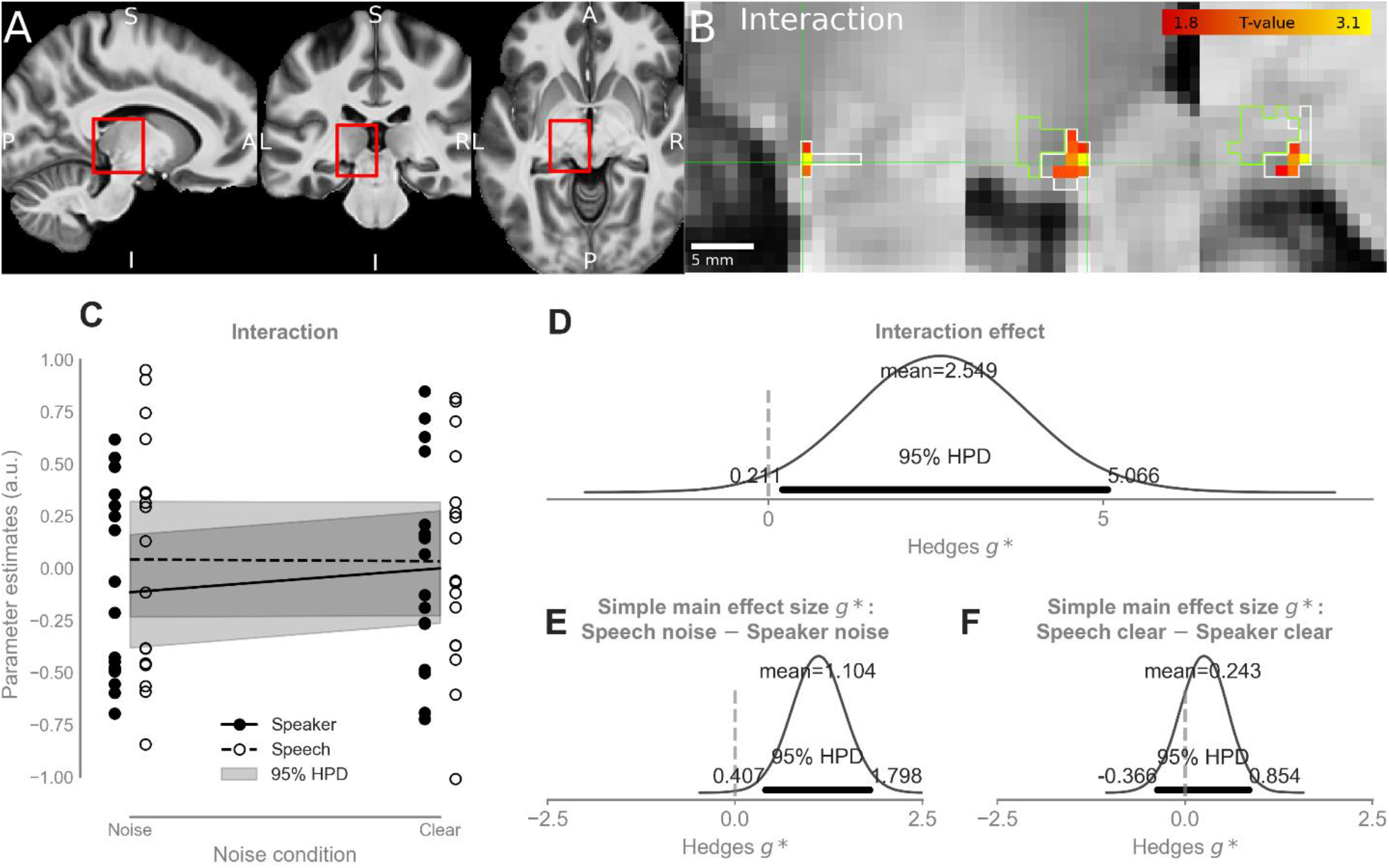
fMRI results. **A.** The mean T1 structural image across participants in MNI space. Red rectangles denote the approximate location of the left MGB and encompass the zoomed-in views in B. Letters indicate anatomical terms of location: A, anterior; P, posterior; S, superior; I, inferior; L, left; R, right. Panels A and B share the same orientation across columns; i.e., from left to right: sagittal, coronal, and axial. **B.** Statistical parametric map of the interaction (yellow-red colour code): (speech task/noise - speaker task/noise) - (speech task/clear - speaker task/clear) overlaid on the mean structural T1 image. Crosshairs point to MNI coordinate (−11, −28, −6). The white outline shows the boundary of the vMGB mask; the green boundary delineates the non-tonotopic parts of the MGB. **C.** Parameter estimates (mean-centred) within the vMGB mask. Open circles denote parameter estimates of the speech task condition; filled circles denote parameter estimates of the speaker task condition. Dashed black line: the relationship between noise condition (noise, clear) and parameter estimates in the speech task. Solid black line: the relationship between noise condition (noise, clear) and parameter estimates in the speaker task. The shaded grey area shows the 95% HPD. **D-F** Bayesian Analysis of the parameter estimates. **D.** The effect size of the interaction: the effect size for the interaction effect was very large (2.549 [0.211, 5.066]) and the HPD excluded zero (indicated by the dashed vertical line). **E**. Simple main effect: speech task/noise - speaker task/noise. The mean effect size was large (1.104 [0.407, 1.798]). The HPD excluded zero. **F**. Simple main effect: speech task/clear - speaker task/clear. The mean effect size was small (0.243 [−0.366, 0.854]). The HPD contained zero.

The results showed that the task-dependent modulation of the left vMGB for the speech task was increased when participants recognised speech - speaker identity in background noise in contrast to speech - speaker identity without background noise (task × noise interaction). This finding cannot be explained by differences in stimulus input as the same stimulus material was used for the speech and the speaker task. The results are also unlikely due to differences in task difficulty between conditions, as the behavioural results showed no detectable differences in performance for the simple main effects.

We did not have a specific hypothesis on the right vMGB, as there is currently no indication that the task-dependent modulation in this region is behavioural relevant (von Kriegstein et al., 2008; Mihai et al., 2019) or dysfunctional in disorders associated with speech-in-noise processing difficulties (Díaz et al., 2012; Tschentscher et al., 2019). Exploring the interaction in the right vMGB revealed no interaction effect as the HPD strongly overlapped zero (g* = −0.544 [−3.093, 2.459]).

##### 3.2.2 Exploratory analyses on the central nucleus of the inferior colliculus (cIC)

In exploratory analyses, we investigated the bilateral cIC involvement during speech processing. The reason for these exploratory analyses were studies using auditory brainstem responses (ABR) during passive listening to speech sounds that have shown that the quality of speech sound representation (i.e., as measured by the frequency following response, FFR) explains inter-individual variability in speech-in-noise recognition abilities (Chandrasekaran et al., 2009; Song et al., 2010; Schoof and Rosen, 2016; Selinger et al., 2016). These findings indicated that there might be subcortical nuclei beyond the MGB that are involved in speech-in-noise perception, potentially also sources in the auditory brainstem, particularly the IC (Chandrasekaran and Kraus, 2010b). Four previous fMRI experiments, however, have shown that there is *no* significant task-dependent modulation (i.e., higher BOLD responses for a speech in contrast to a control task on the same stimuli) of the inferior colliculus (von Kriegstein et al., 2008; Díaz et al., 2012; Mihai et al., 2019). Two of them showed a significant positive correlation between the amount of BOLD response difference between a speech and a control task in the left IC and the speech recognition performance across participants (von Kriegstein et al., 2008, experiment 1 and 2), but the others did not. Thus the role of the IC in speech recognition and speech-in-noise recognition is to date unclear. In the present data, there was a small effect of task in the left cIC (speech - speaker, left g*=0.309 [−0.286, 0.902] and right g*= 0.126 [−0.393, 0.646], however, the HPD overlapped zero. The task × noise interaction contained no explanatory power (left: g*=0.049 [−0.103, 0.202], right: g*=-0.010 [−0.136, 0.111]) and introduced overfitting. We, therefore, excluded it from the model, and the reported results were computed from the model without an interaction term. The correlation between the task-dependent modulation (i.e., speech - speaker task contrast) and the speech recognition scores across participants in the left cIC was not significant in the current study (r=0.44, p=0.074, Figure 7).

**Figure 7.**
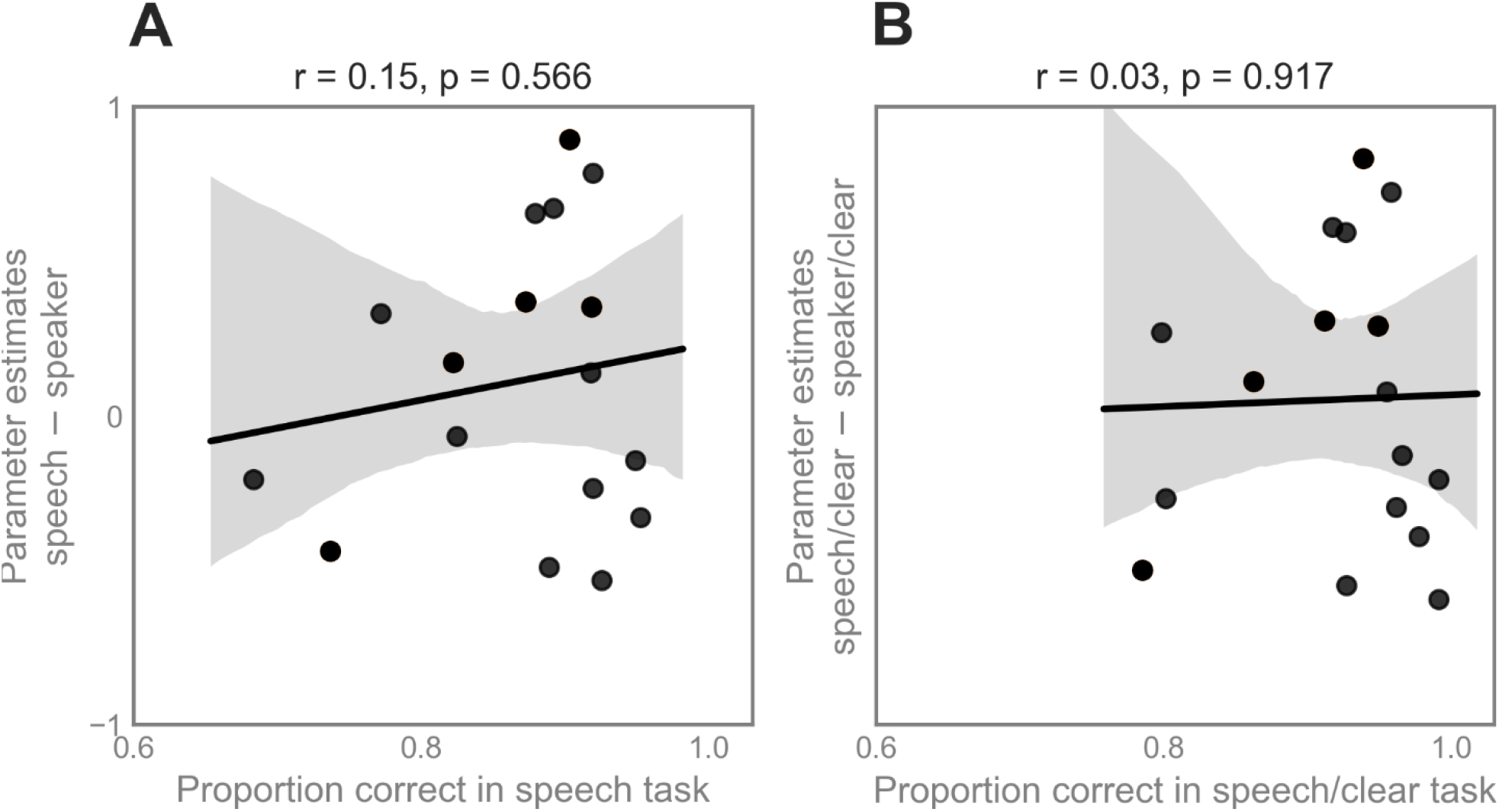
**A** Correlation analysis between the parameter estimates of the contrast Speech – Speaker task in the left cIC and the proportion of hits in the speech task. **B** Correlation analysis between the parameter estimates of the contrast speech/clear – speaker/clear task in the left cIC and the proportion of hits in the speech/clear task. Most data points are close to the ceiling on the right of the behavioural score. For both correlations, the degrees of freedom were 16.

### 4. Discussion

We showed that the task-dependent modulation for speech of the left hemispheric primary sensory thalamus (vMGB) is particularly strong when recognising speech in noisy listening conditions in contrast to conditions where the speech signal is clear. This finding confirmed our a priori hypothesis which was based on explaining speech-in-noise recognition and sensory thalamus function within a Bayesian brain framework. Exploratory analyses showed that there was no influence of noise on the responses for the contrast between speech and speaker task in the right vMGB, or in the auditory midbrain, i.e., the central nuclei of the inferior colliculi (cIC).

Bayesian approaches to brain function propose that the brain uses internal dynamic models to predict the trajectory of the sensory input (Knill and Pouget, 2004; Friston, 2005; Kiebel et al., 2008; Friston and Kiebel, 2009). Thus, slower dynamics of the internal dynamic model (e.g., syllable and word representations) could be encoded by auditory cerebral cortex areas (Giraud et al., 2000; Davis and Johnsrude, 2007; Hickok and Poeppel, 2007; Wang et al., 2008; Mattys et al., 2012; Price, 2012), and provide predictions about the faster dynamics of the input arriving at lower levels of the anatomic hierarchy (Kiebel et al., 2008; von Kriegstein et al., 2008). In this view, dynamic predictions modulate the response properties of the first-order sensory thalamus to optimise the early stages of speech recognition (Mihai et al., 2019). In speech processing, such a mechanism might be especially useful as the signal includes rapid dynamics, that are predictable (e.g., due to co-articulation or learned statistical regularities in words) (Saffran, 2003). In addition, speech often has to be computed online under conditions of (sensory) uncertainty. Uncertainty refers to the limiting reliability of sensory information about the world (Knill and Pouget, 2004). Examples include the density of hair cells in the cochlea that limit frequency resolution, the neural noise-induced at different processing stages, or – as was the case in the current study – background environmental noise that surrounds the stimulus of interest. An internal generative model about the fast sensory dynamics (Knill and Pouget, 2004; Friston, 2005; Kiebel et al., 2008; Friston and Kiebel, 2009) of speech could lead to enhanced stimulus representation in the subcortical sensory pathway and by that provides improved signal quality to the auditory cortex. Such a mechanism would result in more efficient processing when taxing conditions, such as background noise, confront the perceptual system. The interaction between task and noise in the left vMGB is in congruence with such a mechanism. It shows that the task-dependent modulation of the left vMGB is increased in a situation with high sensory uncertainty in contrast to the situation with lower sensory uncertainty. Although the results are in accordance with the Bayesian brain hypothesis, the study was not meant to test directly whether predicticve coding is used in the auditory pathway. To test this it would be necessary to manipulate predictability of the stimuli (Tabas et al., 2020).

Both the speech task and the speaker task required attention to the stimuli. Attention can interact to provide a better decoding of the stimuli we choose to attend to (Schröger et al., 2015), and can optimize predictions of incoming signals (Smout et al., 2019) resulting in a top-down and bottom up signal integration (Gordon et al., 2019). Attention can be formulated in a predictive coding account (Ransom et al., 2017), for example, it could result in increased precision on the prediction. It is to date an open question whether the task-dependent modulation observed for speech recognition in the present and previous studies in sensory thalamic nuclei (von Kriegstein et al., 2008; Díaz et al., 2012, 2018; Mihai et al., 2019) operate through the same mechanisms as attentional modulation (O’Connor et al., 2002; Schneider and Kastner, 2009; Schneider, 2011; Ling et al., 2015)

Speech-in-noise recognition abilities are thought to rely (i) on additional cognitive resources (reviewed in Peelle, 2018) and (ii) on the fidelity of speech sound representation in brainstem nuclei, as measured by auditory brainstem response recordings (reviewed in Anderson and Kraus, 2010). For example, studies investigating speech-in-noise recognition at the level of the cerebral cortex found networks that include areas pertaining to linguistic, attentional, working memory, and motor planning (Salvi et al., 2002; Scott et al., 2004; Bishop and Miller, 2008; Wong et al., 2008). These results suggested that during speech recognition in challenging listening conditions additional cerebral cortex regions are recruited that likely complement the processing of sound in the core speech network (reviewed in Peelle, 2018). The present study showed that besides the additional cerebral cortex region recruitment, a specific part of the sensory pathway is also modulated during speech-in-noise recognition: the left vMGB.

Auditory brainstem response (ABR) recordings during passive listening to speech sounds have shown that the quality of speech sound representation (i.e., as measured by the frequency following response, FFR) explains inter-individual variability in speech-in-noise recognition abilities (Chandrasekaran et al., 2009; Song et al., 2010; Schoof and Rosen, 2016; Selinger et al., 2016) and can be modulated by attention to speech in situations with two competing speech streams (Forte et al., 2017). It is difficult to directly relate the results of these FFR studies on participants with varying speech-in-noise recognition abilities (Chandrasekaran et al., 2009; Song et al., 2010; Schoof and Rosen, 2016; Selinger et al., 2016) to the studies on task-dependent modulation of structures in the subcortical sensory pathway (von Kriegstein et al., 2008; Díaz et al., 2012; Mihai et al., 2019): they involve very different measurement modalities and the FFR studies focus mostly on speech-in-noise perception in passive listening designs. One major candidate for the FFR source is the inferior colliculus. Particularly for speech, the FFR, as recorded by EEG, seems to be dominated by brainstem and auditory nerve sources (reviewed in Chandrasekaran et al., 2014; Bidelman, 2018). The results of the present study, however, do not provide evidence for a specific involvement of the inferior colliculus when recognising speech-in-noise. The choice of syllables for the speech task emphasises predictions at the phonetic level. One possibility is that task-dependent modulation of the left MGB in conditions with high sensory uncertainty, might be particularly relevant for such processing at the phonetic level as the MGB might be optimised for this type of fast-varing information (Giraud et al., 2000; von Kriegstein et al., 2008). Whether the inferior colliculus might play a different role in speech-in-noise processing is an open question.

We speculate that the task-dependent vMGB modulation might be a result of feedback from cerebral cortex areas. The strength of the feedback could be enhanced when speech has to be recognised in background noise. The task-dependent feedback may emanate directly from primary auditory or association cortices, or indirectly via other structures such as the reticular nucleus with its inhibitory connections to the MGB (Rouiller and de Ribaupierre, 1985). Feedback cortico-thalamic projections from layer 6 in A1 to the vMGB, but also from association cortices such as the motion-sensitive planum temporale (Tschentscher et al., 2019), may modulate information ascending through the lemniscal pathway, rather than convey information to the vMGB (Llano and Sherman, 2008; Lee, 2013).

Difficulties in understanding speech-in-noise accompany developmental disorders like autism spectrum disorder, developmental dyslexia, and auditory processing disorders (Alcántara et al., 2004; Chandrasekaran et al., 2009; Wong et al., 2009; Ziegler et al., 2009; Bellis and Bellis, 2015; Schoof and Rosen, 2016; Schelinski and Kriegstein, 2019). In the case of developmental dyslexia, previous studies have found that developmental dyslexics do not have the same amount of task-dependent modulation of the left MGB for speech recognition as controls (Díaz et al., 2012) and also do not display the same context-sensitivity of brainstem responses to speech sounds as typical readers (Chandrasekaran et al., 2009). In addition, diffusion-weighted imaging studies have found reduced structural connections between the MGB and cerebral cortex (i.e., the motion-sensitive planum temporale) of the left hemisphere in developmental dyslexics compared to controls (see Müller-Axt et al., 2017 for similar findings in the visual modality; Tschentscher et al., 2019). These altered structures might account for the difficulties in understanding speech-in-noise in developmental dyslexia. Consider distinguishing speech sounds like “dad” and “had” in a busy marketplace. For typically developed individuals, vMGB responses might be modulated to optimally encode the subtle but predictable spectrotemporal cues that enable the explicit recognition of speech sounds. This modulation would enhance speech recognition. For developmental dyslexics, however, this vMGB modulation may be impaired and may explain their difficulty with speech perception in noise (Boets et al., 2007; Ziegler et al., 2009; Díaz et al., 2012).

In conclusion, the results presented here suggest that the left vMGB is particularly involved in decoding speech as opposed to identifying the speaker if there is background noise. This enhancement may be due to top-down processes that act upon subcortical sensory structures, such as the primary auditory thalamus, to better predict dynamic incoming signals in conditions with high sensory uncertainty.

## Acknowledgements

The study was funded by the European Research Council ERC Consolidator Grant SENSOCOM (647051) and the Max Planck Society.

## Author contributions

PGM: collected data, analysed data, interpreted results, wrote the manuscript, edited the manuscript. NT: conceptualised experiment, programmed experiment, edited the manuscript. KvK: conceptualised experiment, interpreted results, wrote the manuscript, edited the manuscript.

## Notes

Conflict of Interest Statement: The authors declare no competing financial interests.

### Competing Interest Statement

The authors have declared no competing interest.

### Summary of Updates

Improved on text, reorganized images, reformatted.

